# Deep Generative Analysis for Task-Based Functional MRI Experiments

**DOI:** 10.1101/2021.04.04.438365

**Authors:** Daniela de Albuquerque, Jack Goffinet, Rachael Wright, John Pearson

## Abstract

While functional magnetic resonance imaging (fMRI) remains one of the most widespread and important methods in basic and clinical neuroscience, the data it produces—time series of brain volumes—continue to pose daunting analysis challenges. The current standard (“mass univariate”) approach involves constructing a matrix of task regressors, fitting a separate general linear model at each volume pixel (“voxel”), computing test statistics for each model, and correcting for false positives *post hoc* using bootstrap or other resampling methods. Despite its simplicity, this approach has enjoyed great success over the last two decades due to: 1) its ability to produce effect maps highlighting brain regions whose activity significantly correlates with a given variable of interest; and 2) its modeling of experimental effects as separable and thus easily interpretable. However, this approach suffers from several well-known drawbacks, namely: inaccurate assumptions of linearity and noise Gaussianity; a limited ability to capture individual effects and variability; and difficulties in performing proper statistical testing secondary to independently fitting voxels. In this work, we adopt a different approach, modeling entire volumes directly in a manner that increases model flexibility while preserving interpretability. Specifically, we use a generalized additive model (GAM) in which the effects of each regressor remain separable, the product of a spatial map produced by a variational autoencoder and a (potentially nonlinear) gain modeled by a covariate-specific Gaussian Process. The result is a model that yields group-level effect maps comparable or superior to the ones obtained with standard fMRI analysis software while also producing single-subject effect maps capturing individual differences. This suggests that generative models with a decomposable structure might offer a more flexible alternative for the analysis of task-based fMRI data.

## 1. Introduction

For twenty years, functional magnetic resonance imaging (fMRI) has been one of the most prominent experimental modalities in the fields of cognitive and clinical neuroscience, allowing researchers and clinicians to investigate relationships between brain regions and functions or behaviors of interest (e.g., memory consolidation (Fogel et al., 2014; Wittmann et al., 2005), decision making (Hampton and O’Doherty, 2007)). However, the data produced by these experiments — time series of 3D brain images (“brain volumes”) — are both high-dimensional and high noise and continue to pose daunting analysis challenges (Zhang et al., 2020; Burock and Dale, 2000; Zwart et al., 2008). In particular, each brain volume comprises tens to hundreds of thousands of smaller 3D units (voxels) whose blood oxygen level dependent (BOLD) signal (Ogawa et al., 1992) is captured throughout time.

In response to these challenges, numerous studies have sought to model neuroimaging data, most typically focusing on widely available benchmark datasets using resting state connectivity (Ju et al., 2019; Suk et al., 2016; Mao et al., 2019; Tahmassebi et al., 2018), structural (Henschel et al., 2020; Tian et al., 2020; Gunawardena et al., 2017; Zhang, 2018) and fMRI (Gadgil et al., 2020; Riaz et al., 2018; Sarraf and Tofighi, 2016) modalities. However, this large and growing literature, which most often focuses on prediction and classification, has largely ignored *task-based* fMRI, data that result from *designed experiments* aimed at testing particular scientific hypotheses. In particular, such data are generated for purposes of *statistical inference* on covariates of interest. Thus, there is a need for models that address both the complexity of fMRI data and the goals of scientific inference.

Here, we take an alternative approach aimed directly at this problem. Since the quantities of interest to experimenters are spatial effect maps that quantify the effects of covariates on brainwide activity, we use a hybrid approach, combining deep generative models (for the maps) with Gaussian Processes (for each covariate) in a generalized additive model (GAM) approach. That is, we “nest” deep generative models inside a well-understood statistical framework that produces separable (and thus interpretable) effect maps for each covariate of interest. As we show, this model produces results comparable to conventional fMRI analysis methods while better capturing spatial variability and allowing for more flexibility in modeling non-linear effects in data.

### Generalizable Insights about Machine Learning in the Context of Healthcare

As mentioned, task-based fMRI data is a highly popular imaging modality in both basic and clinical neuroscience, as it allows researchers to test hypotheses about brain function in health and disease states through carefully designed and controlled experiments. For example, classic tasks like the *n-back paradigm* (Koshino et al., 2005; Ragland et al., 2002; Blokland et al., 2008) have allowed clinical researchers to discover bio-markers of working memory deficits typically observed in psychiatric diseases like schizophrenia (Jansma et al., 2004; Callicott et al., 2003; Nicodemus et al., 2010) and to relate these to variables of interest such as clinical severity scores (Hashimoto et al., 2009) and genotypes for markers involved in dopaminergic or glutamatergic neurotransmission (Egan et al., 2001; Apud et al., 2017; Egan et al., 2004). As another example, fMRI studies have shown that children diagnosed with ASD fail to activate sound-processing brain regions (e.g., STS) in response to vocal sounds (Gervais et al., 2004) and have overall altered activity patterns in key brain regions associated with social functioning (Philip et al., 2012), which might explain the social deficits typically observed in ASD. fMRI has also helped unveil abnormalities in fronto-limbic activation patterns (e.g., reduced IFG activity) which has been linked with episodes of mania typically seen in bipolar disorder (Chen et al., 2011). Though these are only a a few examples, broadly, task-based fMRI is widely used as both a diagnostic and a research method across the brain sciences, where it serves as a key link between higher-level cognitive function, brain spatio-temporal dynamics and putative molecular and genetic mechanisms of disease.

The alternative fMRI analysis framework we propose here compares favorably to the current state of the art detailed below while harnessing the flexibility of deep generative models to capture high-dimensional and complex data. Moreover, this approach retains the interpretable constructs (effect maps) with which neuroscientists and clinicians are familiar, allowing it to serve as a bridge between established methods and modern techniques. Ultimately, developing better methods to model these data will empower clinicians and neuroscientists to utilize fMRI to its full potential and discover valuable links between molecular mechanisms, neural function and high-level behavior in health and disease.

## 2. Related Work

Standard approaches to fMRI data analysis routinely make use of compression and dimension reduction-based approaches such as independent component analysis (ICA) (Bai et al., 2007; Calhoun et al., 2009), canonical correlation analysis (Friman et al., 2001; Hardoon et al., 2007; Lin et al., 2014), and less frequently sparse dictionary learning (Lee et al., 2011; Eavani et al., 2012; Lv et al., 2015), but these approaches typically rely on strong assumptions like linearity and spatial independence which are violated for fMRI data. In addition, SVM-based classification methods, known within the field under the name multivoxel pattern analysis (MVPA) (Norman et al., 2006; Mahmoudi et al., 2012) have been widely used to localize particular kinds of task effects within the brain while making weaker statistical assumptions.

More recently, there has been an explosion of work using deep networks to model different modalities of brain imaging data, including resting state (rs-fMRI), structural, and diffusion tensor imaging (DTI). For example, deep neural nets have been used to extract latent features from fMRI data, which can then be used for downstream classification tasks (Huang et al., 2016; Jang et al., 2017; Suk et al., 2015; Han et al., 2015). They have also been used for de-noising neuroimaging data (Yang et al., 2020; Zhao et al., 2020). These models have also been extensively used to classify imaging data into different diagnostic groups or into different brain networks (Khosla et al., 2019; Li et al., 2018; Sarraf and Tofighi, 2016; Ju et al., 2019; Ren et al., 2017) and to predict values of regressors of interest from imaging data (Chen et al., 2019; Jiang et al., 2020; Jonsson et al., 2019). These applications have been primarily driven by the advent of public datasets focused on diagnosis and disease prediction, along with the wide availability of modalities like rs-fMRI and DTI, which are comparatively easier to obtain for large cohorts.

However, markedly less attention has been given to the use of deep unsupervised approaches (e.g., VAEs) in capturing high-level representations of brain network organization and dynamics directly from low-level brain imaging data. As an example, Huang et al. (2017) utilized a deep convolutional autoencoder to extract high-level representations from task-based fMRI data and found that such representations not only have good correspondence with theoretical models of brain response but are also superior to dictionary learning approaches in detecting task-related regions. Suk et al. (2016) coupled a deep Auto-Encoder with a hidden Markov Model (HMM) to learn non-linear functional relations among brain regions and estimate the dynamics of such relations from rs-fMRI data. More recently, Qiang et al. (2020) adopted a neural architecture search (NAS), along with a deep belief network (DBN), to achieve an optimal scheme for modeling task-specific and resting state functional brain networks in an unsupervised fashion. Moreover, some of these studies (Matsubara et al., 2021, 2018) have utilized unsupervised approaches to differentiate between diagnostic criteria, and the high levels of individual variability typically observed in fMRI data.

The most closely related work to our model is that of Zabihi et al. (2021), which analyzed task-based data using a standard VAE. They visualized latent representations across different tasks, showing that these were distinguishable, but they did not examine the role of task covariates or experimental designs. Likewise, Zhao et al. (2019) used a VAE to analyze MRI data with age considered as a covariate, but they did not model task-based data. Thus, to our knowledge, ours is the first work to use autoencoders to address statistical inference in designed experiments, the major goal of scientific analysis in functional MRI experiments. Moreover, the structure of our model differs from the standard VAE in producing covariate-specific, interpretable effect maps for variables of interest, similar to those produced by standard GLM-based approaches.

## 3. Methods

### 3.1. Mass univariate regression for fMRI analysis

We denote the brain BOLD signal at time *t* as ***x***_*t*_, with individual voxel intensities *x_tj_*, where we let *j* range over all spatial locations. We further assume a set of covariates defined at each time, {*c_tα_*}, where *α* indexes not only experimental conditions but also nuisance variables like head motion and respiration for which we would like to control. In the mass univariate approach, the effects of these regressors are modeled independently for each voxel:

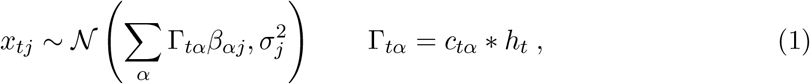

where **Γ** is the experimental design matrix, each column of which is formed by convolving a particular regressor with the hemodynamic response function *h_t_*, the (assumed) transfer function between local brain activity and BOLD response. The *β_αj_* estimated from data can be viewed as effect *maps* (one per regressor *α*) as a function of spatial location *j*. In practice, after voxelwise model fitting, summary statistics and estimated covariances are aggregated across separate experimental runs and participants in a manner equivalent to mixed effects models (Beckmann et al., 2003; Woolrich et al., 2004). The downside of this approach, mentioned above, is that there is no sharing of statistical strength across voxels in estimating the maps *β_αj_*. Thus, while cluster corrections and related methods successfully control for false positives, they only indirectly control for false negatives, typically by setting liberal voxelwise statistical thresholds before resampling. In addition these methods ignore the natural correlation structure, both spatial and temporal, present in the data.

### 3.2. The GAM-VAE model

In contrast, we propose to model entire brain volumes using a single generative model based on variational autoencoders (VAEs) (Kingma and Welling, 2013; Rezende et al., 2014). In the VAE, one assumes a generative model in which the data, ***x***_*t*_, are drawn from a distribution *p_θ_*(**x**|**z**) that depends on a lower-dimensional latent variable **z** and is parametrized by ***θ***. Inference for **z** then proceeds by choosing a class of posterior distributions *q_ϕ_*(**z**|**x**) parametrized by ***ϕ*** and minimizing the Kullback-Leibler divergence *D_KL_*(*q_ϕ_*||*p_ϕ_*) over (***θ, ϕ***). More concretely, we take

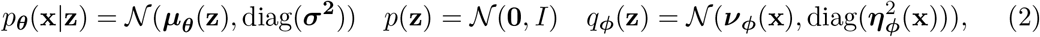

where ***μ***, ***v***, and ***η*** are functions approximated by deep neural networks, *I* is the identity matrix, and we allow a separate variance 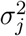 for each voxel. We then maximize a stochastic approximation to the evidence lower bound (ELBO):

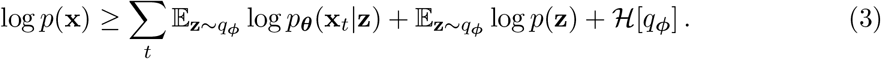

Here 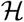 is the differential entropy and the expectation is approximated by drawing samples (Blei et al., 2017). We have also assumed that observations ***x***_*t*_ are independent and identically distributed, though this is known to be a poor approximation in the case of highly temporally autocorrelated fMRI data. We justify this on three grounds: First, this mirrors standard fMRI analysis, in which autocorrelation is assumed to be adequately modeled by the autocorrelation of regressors and the convolution by *h_t_* in (1). Second, the sluggishness of the BOLD response in relation to underlying neural activity argues for experimental designs in which temporal dynamics are treated as nuisance variables to be averaged over. Finally, as we shall see, our model naturally lends itself to extensions where the evolution of the latent variables ***z***_*t*_ can be modeled explicitly as in, e.g., (Le et al., 2017; Maddison et al., 2017; Naesseth et al., 2018) or using recurrent neural networks, though we do not pursue that here.

Of course, (2) does not include the effects of the covariates **c**. The most general extension is to include these values as inputs to the maps ***μ***, ***v***, and ***η***, but in the case of ***μ***, this creates difficulties for interpretability, since the effect of *c_α_* then potentially depends on the values of all other covariates. Instead, we indeed take ***v***(**x**, **c**), and ***η***(**x**, **c**) but assume for ***μ*** a generalized additive model (GAM) (Hastie and Tibshirani, 1990) in which the observed signal is a sum of covariate-specific effects:

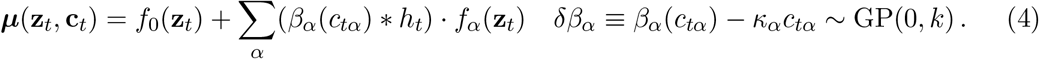

That is, we assume a set of effect maps *f_α_* parameterized by neural networks and a set of covariate-dependent gain functions *β_α_* whose nonlinearities are drawn from a Gaussian Process with kernel *k* (Williams and Rasmussen, 2006). As in the general linear approach, each covariate effect *β*, representing the neural response, is convolved with the hemodynamic response *h_t_*, and this result is used to scale the effect map *f_α_*. Note that the Gaussian process in (4) does not model *β* itself but the *difference* between the covariate effect and its best linear approximation, *δβ*. While the GP is theoretically flexible enough to model *β* itself, we found that in practice the residual formulation performed better, perhaps because of the difficulty of the GP in capturing a pure linear trend. As a result, the linear coefficients *κ_α_* represent the presence or absence of a linear covariate effect, while credible intervals for the GP allow us to assess potential nonlinearity. Of course, since a nonzero *κ_α_* is not identifiable (it can be absorbed into the definition of *f_α_* provided the GP residual is also rescaled), we often choose *κ_α_* ≡ 1 when it does not vanish. The result is a model that incorporates the natural covariance structure of brain volumes via the VAE, potentially nonlinear covariate responses via the Gaussian Process, and does so in a manner that preserves experimental interpretability (Figure 1).

**Figure 1:**
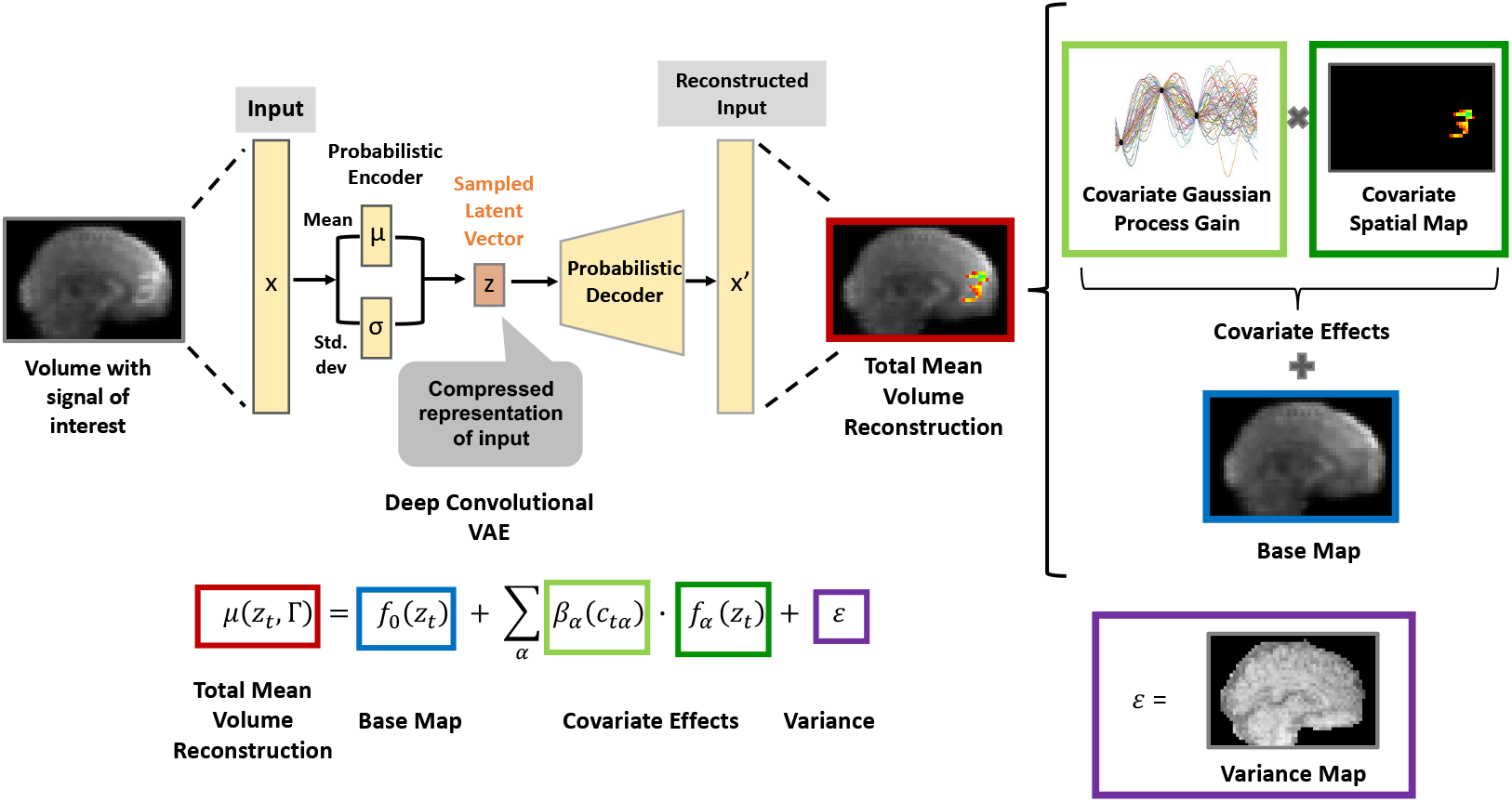
Schematic of the GAM-VAE fMRI model. Each fMRI volume is compressed to a lower-dimensional representation using a probabilistic encoder network. Latent space vectors are then sampled and fed to a probabilistic decoder, which yields a base map (blue) and a set of covariate-specific maps (dark green). The gain of each covariate map is the sum of a linear effect term (*κ_α_ct_α_*) and a potential non-linearity modeled by a Gaussian process (light green). Finally, the full mean reconstruction of the input is obtained by adding the base and the scaled covariate maps. Variance is modeled on a per-voxel basis, yielding a separate variance map (purple).

### 3.3. Network Architecture, Training and Map Reconstruction

For the encoder and decoder neural networks, we used a standard convolutional neural network architecture previously used for images (Goffinet et al., 2019), generalized to 3D convolutions for brain volumes. The encoder consisted of 5 convolutional layers, 3 batch normalization layers, and 4 fully-connected layers (see Appendix **A** for details). Networks responsible for the mean (*f*_0_), covariate-specific (*f_α_*) effects, and for the lower triangular and diagonal matrices used to construct the Cholesky factor of the covariance matrix shared both convolutional and fully-connected early layers and so shared feature sets. Analogously, the decoder network consisted of 4 fully-connected layers, followed by 5 transposed 3D- convolutional layers and 3 batch-normalization layers (Appendix **A**).

For the covariate-specific Gaussian Processes (4), we used the sparse variational approximation of (Hensman et al., 2015). That is, we assume an approximate Gaussian Process posterior, *q*(*δβ*), defined at a small number of inducing point locations, and we optimize the parameters of this distribution (see Appendix **B** for details). More specifically, we used a small number of uniformly spaced inducing points per covariate and a radial basis function (Gaussian) kernel whose length scale and variance were jointly trained with the VAE. For our experiments, rather than perform full GP inference, we instead used a maximum likelihood approach that optimized the mean and kernel parameters of the GP prior (Appendix **B**):

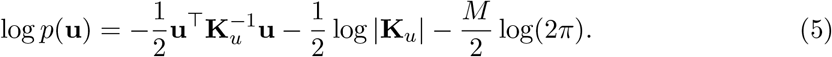

Here, **u** is the vector of values of a single GP at the inducing points, **K**_*u*_ is the prior covariance derived from the kernel, and *M* = 6 is the number of inducing points.

The full GAM-VAE model was trained via using the Adam optimizer, approximating gradients using samples from the posterior *q_ϕ_*(**z**_*t*_|**x**_*t*_) to compute expectations in (3) (Blei et al., 2017). Optimization attempted to minimize the value of a composite objective,

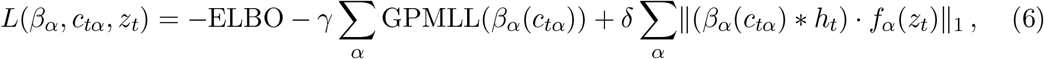

which consisted of the negative ELBO (3), the (weighted) negative marginal log likelihood of the GP (5), and an L_1_ regularization term that encouraged sparse effect maps. Hyperparameters *γ* and *δ* were chosen by grid-search over different possible values sets ((1, 10, 50, 100) for *γ* and (0, 0.005, 0.05 and 0.1) for *δ*, respectively) to yield mean reconstructions with the highest visual quality (see Appendix **C** for results obtained using different hyperparameter values). For our synthetic signal experiments (see below), we used (*γ, δ*) = (10, 0) and for the experiments looking into the biological, V1 effect (also listed below) we used (*γ, δ*) = (10,0.005).

## 4. Cohort and Data

Data consisted of 6 subjects (5 female, age = 33.2 ± 7.9 yrs, 5 right-handed), whose fMRI data were acquired as a part of the neurofeedback task of the NKI-Rockland sample (Nooner et al., 2012). Briefly, the NKI-Rockland sample consists of data from a total of 180 residents of Rockland, Westchester, or Orange Counties (NY) or Bergen County (NJ), aged 21–45 yrs. This sample uses *minimal* psychiatric exclusion criteria — i.e., it excludes subjects with Global Assessment of Function (GAF; DSM-IV) below 50, history of acute or chronic substance abuse, of psychiatric hospitalization, diagnosis of schizophrenia or prior suicide attempts requiring medical intervention. It also excludes individuals with other chronic or serious medical conditions (e.g., epilepsy, TBI, stroke). For a full description of medical exclusion criteria, see McDonald et al. (2017).

Data were acquired using a 3T Siemens Magnetom TrioTrim scanner (TR = 1400 ms, TE = 30 ms, voxel size = (2.0mm)^3^, flip angle = 65 degrees, FoV = 224 mm) (Nooner et al., 2012). The Checkerboard Task, in which subjects were presented either a checkerboard or a fixation cross on a gray background screen, utilized a block-design structure, with each block lasting 20s and a total of four blocks per category (i.e., four checker, four fixation-cross). This task is known to evoke strong activity in brain areas responsible for visual processing (Nooner et al., 2012).

### 4.1. Data Preprocessing

For each subject, data preprocessing consisted of motion-correction, registration to each subject’s structural scan (T1w), warping to MNI space, brain-masking and down-sampling to 41 × 49 × 35. No spatial smoothing was applied, and slice-timing correction was omitted, since data are multi-band. Finally, single volumes were normalized globally (i.e., across all subjects) prior to being fed to the GAM-VAE model. Task covariates included the task period itself (representing the effect of checkerboard presentation) plus 6 motion regressors (3 translational, 3 rotational). For analysis, only brain activity-related covariates (in this case task period) were convolved with the hemodynamic response function (HRF) in (4).

### 4.2. Evaluation Approach/Study Design

The goal of our experiments is to compare the GAM-VAE approach to the current standard of practice analysis approach, the mass univariate model. For these analyses, we used FSL (Jenkinson et al., 2012; Woolrich et al., 2009; Smith et al., 2004). We also compared selected analyses with another widely used software package, SPM (Penny et al., 2006), which produced very similar results. Since standard methods have been the subject of nearly two decades of active development, our tests were aimed at providing evidence that (a) our model, which uses approximate Bayesian methods, nonetheless controls for false positive rates when a ground truth effect is known to be present; (b) these properties degrade gracefully as the signal to noise ratio of the effect is lowered; and (c) the GAM-VAE could produce equivalent (or better) effect maps to those found via standard approaches. That is, we investigate both the power and, to a limited extent, the calibration of our model, as well as its interpretability.

### 4.3. Results on Synthetic Experiments

A key difficulty in assessing the performance of statistical methods on fMRI data is that true synthetic data are challenging to simulate, while ground truth effects are unknown in real data. Thus, to test how well the proposed model can recover a known ground truth signal, we added a synthetic regressor, a large (13 × 13) hand-written “3,” to the checker dataset volumes, creating new, altered datasets. More specifically, the added signal was placed at a constant location in the frontal lobe, with varying signal intensities (2000, 1500, 1000, and 400 arbitrary units (a.u.)). The intensity of the added signal was constant across all voxels and varied only across tests and was active only during control/fixation blocks so as to overlap minimally with time points at which the visual checkerboard stimulus was presented.

We trained the GAM-VAE model on these altered datasets for 400 epochs, at which point convergence was achieved and reconstructions had good visual quality on inspection. For these control simulations, we randomly initialized all contrast maps. Additionally, since the added signal is artificially introduced (and, therefore, not subject to hemodynamic filtering), we did not convolve the GP for the synthetic regressor with the HRF.

Figure 2 shows the resulting average maps (across-participants) for the synthetic regressor effect. Panel A shows a map of the ground-truth signal, overlaid on an anatomical standard template. Panel B shows the average maps generated by our model capturing the synthetic signal across four different signal intensities (e.g., 400, 1000, 1500 and 2000 a.u.). For each intensity, the model was trained using 3 different seeds (rows in panel B). As expected, the model can correctly recover the shape of the synthetic effect for higher signal intensity values, with reconstruction degrading as signal strength decreases.

**Figure 2:**
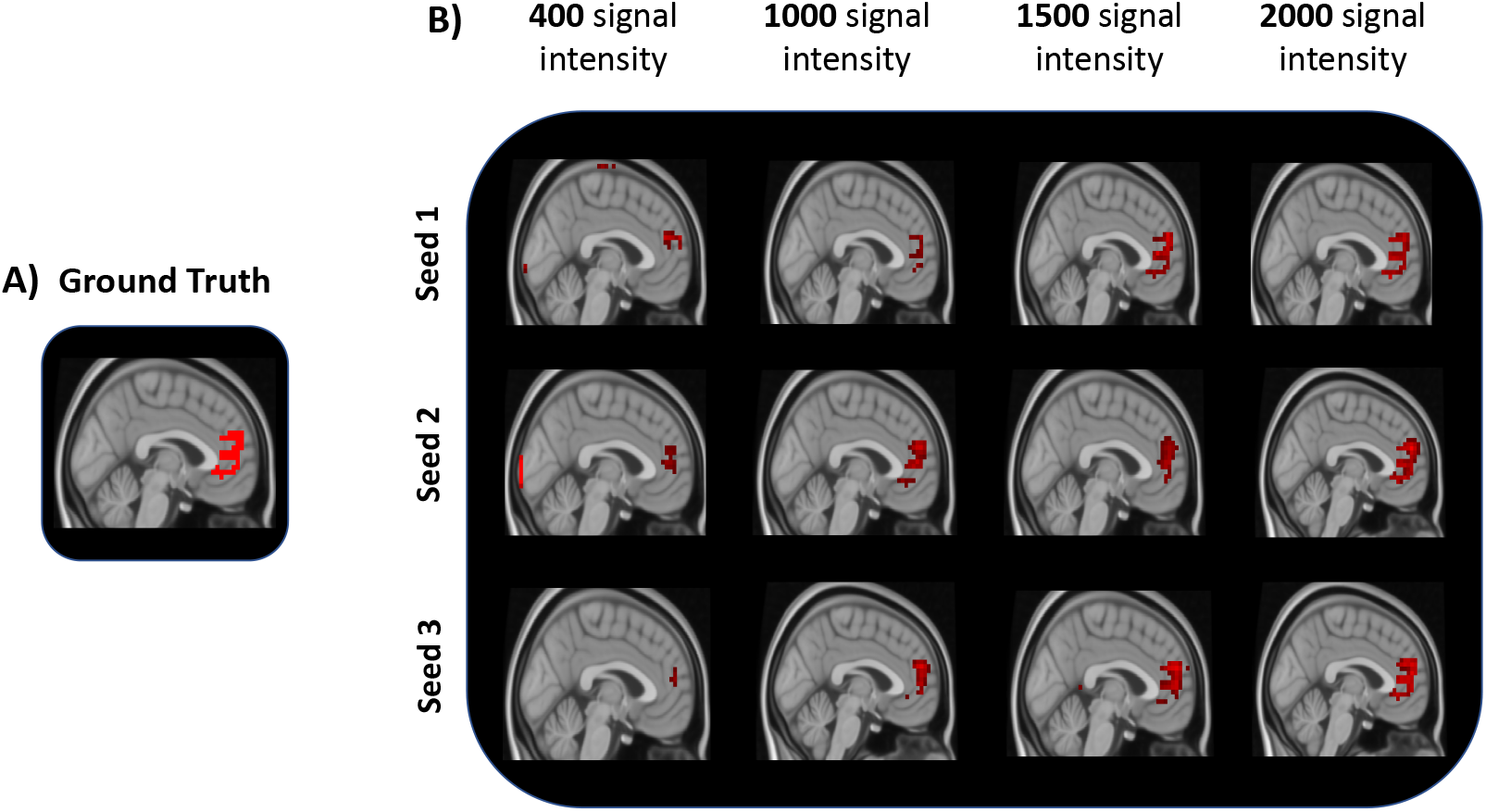
Performance of the GAM-VAE model on synthetic data. **A**: Ground truth effect map for synthetic regressor added. **B**: Average synthetic regressor maps at different signal intensity values (3 different seeds per intensity).

Figure 3 shows a quantitative assessment of our model’s ability to estimate the effect size of this synthetic regressor. To perform this analysis, we mimicked a real experiment (in which the shape of the signal would be unknown) by analyzing data within a spherical mask centered on the location of the synthetic signal. We then defined total effect size as the sum of the intensities of all voxels within the mask. Likewise, we performed standard mass univariate GLM analysis on these data and used the same masking approach (mask was applied to the un-thresholded and un-corrected average contrast maps), and compared both results to the ones obtained for the ground truth maps. As Figure 3 shows, both our proposed GAM-VAE model and the GLM underestimate effect sizes in general, though our model comes closer to correctly estimating effect sizes. In fact, for the smaller intensities (400, 1000) our estimate lies within 1 standard deviation of ground-truth.

**Figure 3:**
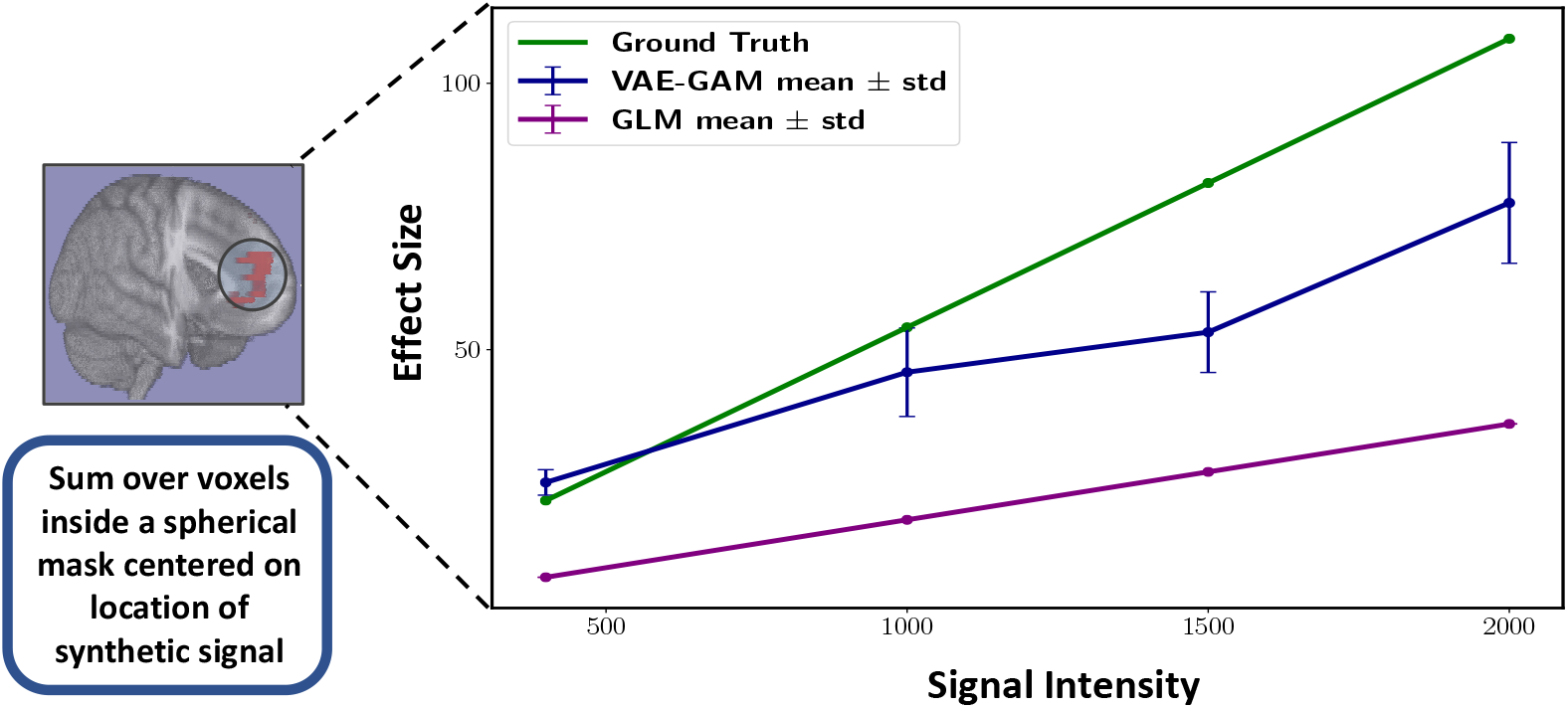
Effect Sizes predicted by GAM-VAE model and GLM vs. ground truth for different signal intensities. Effect sizes were computed as the sum of surviving voxels inside a spherical mask centered on location of the original groundtruth signal. Green, blue and purple lines represent ground-truth, GAM-VAE and GLM effect size estimates, respectively. Error bars represent standard deviation across three independent simulations carried out for each signal strength. Both models underestimate the ground truth effect size, though the GAM-VAE is more accurate, especially for lower signal strengths.

## 5. Results on Visual Stimulation Data

To assess the ability of the GAM-VAE to infer well-validated biological effects, we trained the model using a benchmark dataset consisting of repeated presentations of a visual stimulus. For these data, the model was trained for 400 epochs, at which point convergence was achieved and reconstructions had good quality upon inspection. To aid convergence, we initialized the visual task effect map using a scaled average of the task maps generated using the standard GLM approach.

Figure 4 shows the resulting group average task contrast map generated by our model, along with the average contrast map obtained using the standard GLM approach. Note that the GAM-VAE effect map appears smoother and exhibits both fewer spurious activations outside of visual cortex and fewer “missing” voxels than the GLM map. In Figure 5 we also show comparable task contrast (visual stimulus on versus off) maps for four sample participants. These maps are generated by averaging across all task-containing volume reconstructions for each given subject. As can be seen, the exact location and spatial extent of the stimulus effect varies slightly from subject to subject, though all subjects show a consistent response around visual area V1 (see maps for the remaining 2 subjects in Appendix **D**). Thus, our model is able to capture both population-level inferences and individualized effect estimates.

**Figure 4:**
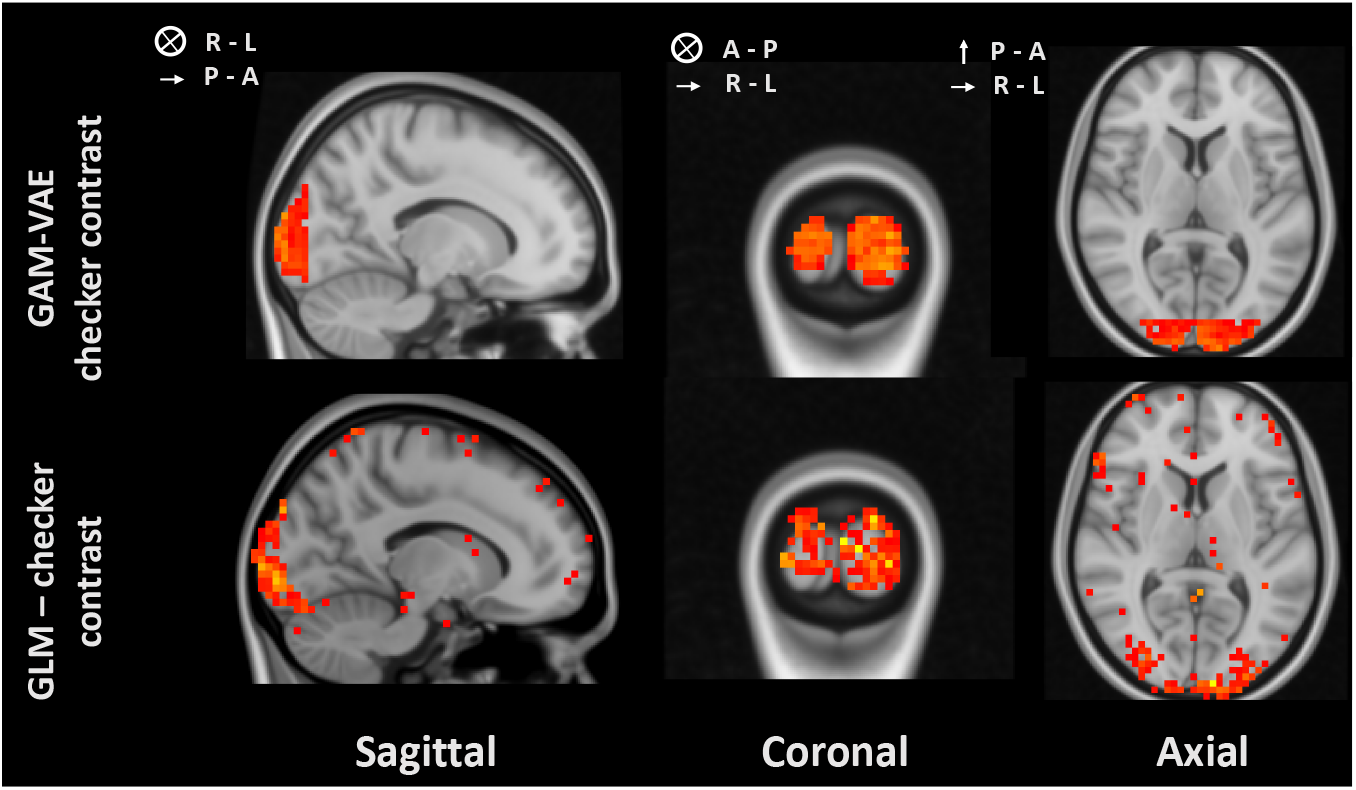
Average effect maps for visual data set experiments. Group average maps generated by the proposed GAM-VAE model (top row) vs. the GLM (bottom row). Note that GAM-VAE model not only captures V1 effect appropriately, but also produces smoother and more contiguous clusters, with less spurious activations outside of V1.

**Figure 5:**
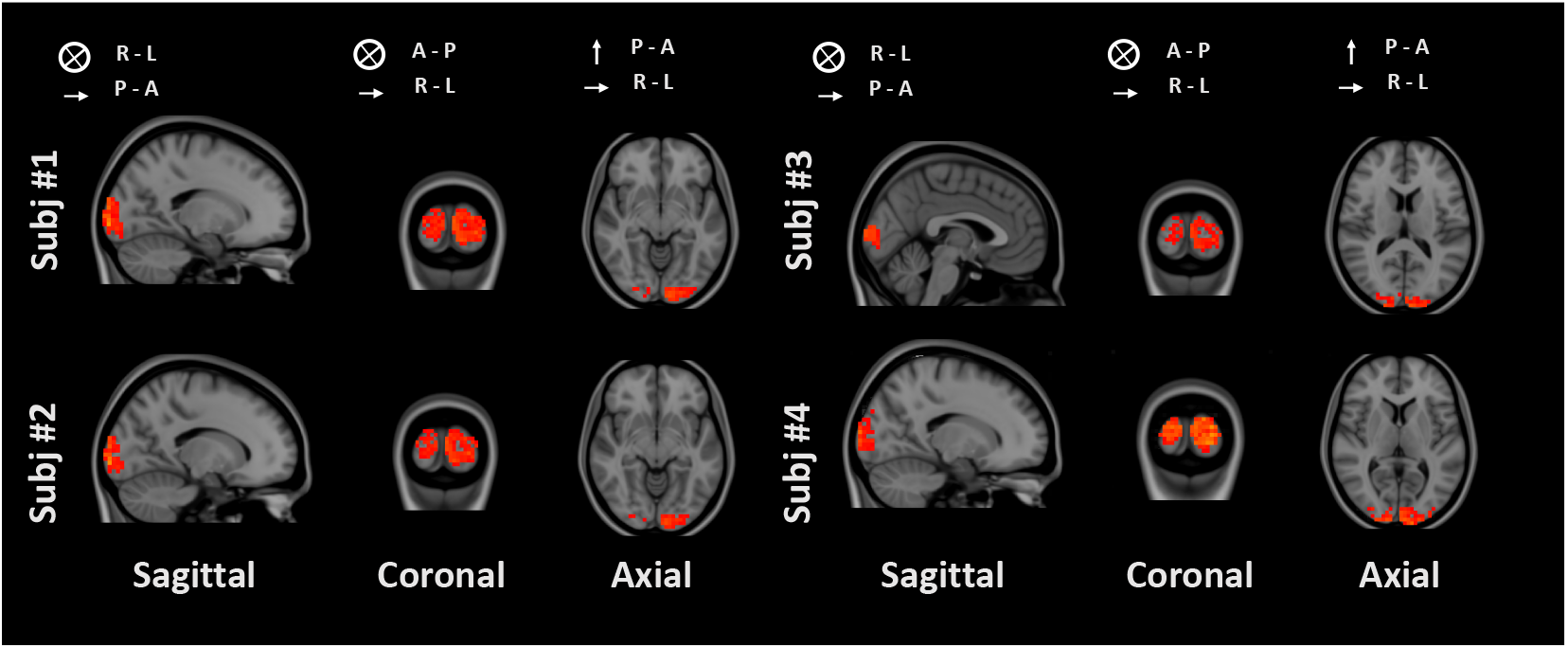
Single subject task effect maps for 4 sample participants. Each map was generated by averaging over all task-containing volume reconstructions for a given subject. Note that the exact location and spatial extent of inferred task covariate effects varies slightly from subject to subject. Maps are displayed at the location at which the main activation cluster is most easily visible for a given subject.

It is also of interest to note that analysis run-time across all levels took approximately 8 hours total for this small cohort (*n* = 5). While this is comparable to run times using the standard approach, our model has the potential to yield significantly lower run times for larger data sets (e.g., data sets with more runs and more subjects). This is because our model does not require running separate analyses for each level separately (e.g., each run, each subject, and then across subjects) before aggregating results and, additionally, it does not require the expensive re-shuffling procedures necessary for *post hoc* inflated false positive control (e.g., for searchlight multi-voxel pattern analysis (Allefeld and Haynes, 2014)).

## 6. Discussion

In this work, we have proposed a new framework for modeling and analyzing task-based fMRI data using deep generative models. More specifically, we nest a deep convolutional auto-encoder inside a GAM framework so as to produced separate and interpretable covariate effect maps. These effect maps are in turn scaled by covariate-specific gains modeled as the sum of a best linear effect estimate and a potential non-linearity (modeled by covariate-specific, one-dimensional Gaussian processes). This approach not only maintains the highly desirable properties of existing fMRI analysis methods (effect separability, generation of spatial effect maps) but also provides more flexibility (e.g., potential non-linear gains) and avoids known drawbacks encountered with the standard GLM approach. For instance, the VAE-GAM framework proposed here models entire volumes at once, which allows it to better account for the inherent spatial auto-correlations observed in fMRI data and avoids some of the statistical testing challenges like inflated false positive rates that require careful correction in the mass-univariate approach.

As we have shown, the proposed VAE-GAM model allows us to correctly recover the shape, size and location of a known synthetic signal added to BOLD data. Additionally, it is capable of producing higher-quality effect maps for true biological signals of interest. In particular, our maps exhibit smoother effect clusters, fewer spurious voxels outside of the main effect cluster, and effect maps for individual participants.

Taken together, the results presented here suggest that deep generative models might provide a new approach to analyzing fMRI data while accounting for some of the modeling challenges inherent to this imaging modality. Ultimately, a more flexible fMRI analysis approach based on a reduced dimensionality latent variable might better model effects of interest to neuroscientists and clinicians, particularly patterns of structured *spatiotemporal* activity associated with cognitive functions and disease processes. Therefore, developing, validating, and perfecting models like this one should be of great value to *both* basic and clinical neuroscience research communities.

### Limitations

Here, we do not directly attempt to incorporate the temporal autocorrelation inherent in fMRI data into our generative structure. Instead, we model this time dependency indirectly using a convolution operation between the stimulus function and the hemodynamic response curve, in a similar manner to what is typically done in a standard GLM analysis. Although we do not directly tackle this issue here, modeling a time series of volumes directly and generating a corresponding sequence of latent variables (Naesseth et al., 2018; Le et al., 2017; Maddison et al., 2017) will yield an even more flexible modeling framework, capable of accounting for both spatial and temporal effect variations which is not achieved by any fMRI analysis method currently in use.

Another limitation lies in the fact that VAEs are known to underestimate posterior variance and, therefore, might yield overconfident results (Böhm et al., 2019). Addressing this issue and arriving at a model capable of providing strong statistical guarantees to researchers interested in quantifying and comparing effect sizes across subjects and conditions is also an important area for improvement. Developing such tools would also be broadly interesting and relevant to the ML community, as it would address a major problem encountered in calibrating VAE-based models.

Finally, the average effect maps showcased here for the checker data set were obtained by initializing the GAM-VAE model with an average map computed using the standard GLM framework. Although our model is capable of generating an effect map with a cluster on the correct location with random initialization alone, we found that providing a better initialization significantly improved the quality of the final map. Therefore, one of limitations of this method lies in the fact that it does rely on non-trivial initializations to produce high-quality results for real, biological effects. Future iterations of this work should attempt to make model more robust to initialization choices.

## Acknowledgements

This work was supported by the National Institutes of Health (R01MH124112).

## References

Carsten Allefeld and John-Dylan Haynes. Searchlight-based multi-voxel pattern analysis of fmri by cross-validated manova. NeuroImage, 89:345–357, 2014. ISSN 1053-8119. doi: 10.1016/j.neuroimage.2013.11.043.

J. Apud, Y. Tong, R. Rasetti, T. Vargas, J. Callicott, D. Dickinson, D. Weinberger, V.S. Mattay, and K.F. Berman. Cortical information processing in patients with schizophrenia is modulated by tolcapone: Role of comt val158met genotype. Schizophrenia Bulletin, 43, 2017.

B. Bai, P. Kantor, A. Shokoufandeh, and D. Silver. fmri brain image retrieval based on ica components. In Eighth Mexican International Conference on Current Trends in Computer Science (ENC 2007), page 10–17, Sep 2007. doi: 10.1109/ENC.2007.32.

Christian F Beckmann, Mark Jenkinson, and Stephen M Smith. General multilevel linear modeling for group analysis in fmri. Neuroimage, 20(2):1052–1063, 2003.

David M Blei, Alp Kucukelbir, and Jon D McAuliffe. Variational inference: A review for statisticians. Journal of the American statistical Association, 112(518):859–877, 2017.

Gabriëlla A.M. Blokland, Katie L. McMahon, Jan Hoffman, Gu Zhu, Matthew Meredith, Nicholas G. Martin, Paul M. Thompson, Greig I. de Zubicaray, and Margaret J. Wright. Quantifying the heritability of task-related brain activation and performance during the n-back working memory task: A twin fmri study”, journal = “biological psychology. 79 (1):70 – 79, 2008. Genetics and Imaging in Neuroscience.

M. A. Burock and A. M. Dale. Estimation and detection of event-related fmri signals with temporally correlated noise: A statistically efficient and unbiased approach. Human Brain Mapping, 11:249–260, 2000.

Vanessa Böhm, François Lanusse, and Uroš Seljak. Uncertainty quantification with generative models. 2019. URL https://arxiv.org/abs/1910.10046v1.

Vince D. Calhoun, Jingyu Liu, and Tulay Adali. A review of group ica for fmri data and ica for joint inference of imaging, genetic, and erp data. NeuroImage, 45(1, Supplement 1):S163–S172, 2009. ISSN 1053-8119. doi: 10.1016/j.neuroimage.2008.10.057.

J.H. Callicott, M.F. Egan, V.S. Mattay, A. Bertolino, A.D. Bone, B. Verchinski, and D.R. Weinberger. Abnormal fmri response of the dorsolateral prefrontal cortex in cognitively intact siblings of patients with schizophrenia. American Journal of Psychiatry, 160:709–719, 2003.

Chang-Le Chen, Yao-Chia Shih, Horng-Huei Liou, Yung-Chin Hsu, Fa-Hsuan Lin, and Wen-Yih Isaac Tseng. Premature white matter aging in patients with right mesial temporal lobe epilepsy: A machine learning approach based on diffusion mri data. NeuroImage: Clinical, 24:102033, 2019. ISSN 2213-1582. doi: 10.1016/j.nicl.2019.102033.

Chi-Hua Chen, John Suckling, Belinda R. Lennox, Cinly Ooi, and Ed T. Bullmore. A quantitative meta-analysis of fmri studies in bipolar disorder. Bipolar Disorders, 13(1): 1–15, 2011. ISSN 1399-5618. doi: https://doi.org/10.1111/j.1399-5618.2011.00893.x.

H. Eavani, R. Filipovych, C. Davatzikos, T. D. Satterthwaite, R. E. Gur, and R. C. Gur. Sparse dictionary learning of resting state fmri networks. In 2012 Second International Workshop on Pattern Recognition in NeuroImaging, page 73–76, 2012. doi: 10.1109/PRNI.2012.25.

M.F. Egan, T.E. Goldberg, B.S. Kolachana, J.H. Callicott, C.M. Mazzanti, R.E. Straub, D. Goldman, and D.R. Weinberger. Effect of COMT Val^108/158^ Met genotype on frontal lobe function and risk for schizophrenia. Proceedings of the National Academy of Sciences, 98, 2001.

M.F. Egan, R.E. Straub, T.E. Goldberg, I. Yakub, J.H. Callicott, A.R. Hariri, V.S. Mattay, A. Bertolino, T.M. Hyde, C. Shannon-Weickert, M. Akil, J. Crook, R.K. Vakkalanka, R. Balkissoon, R.A. Gibbs, J.E. Kleinman, and D.R. Weinberger. Variation in grm3 affects cognition, prefrontal glutamate, and risk for schizophrenia. Proceedings of the National Academy of Sciences, 101, 2004.

S. M. Fogel, G. Albouy, C. Vien, R. Popovicci, B. R. King, R. Hoge, S. Jbabdi, H. Benali, A. Karni, P. Maquet, J. Carrier, and J. Doyon. Fmri and sleep correlates of the age-related impairment in motor memory consolidation. Human Brain Mapping, 35:3625–3645, 2014.

Ola Friman, Jonny Cedefamn, Peter Lundberg, Magnus Borga, and Hans Knutsson. Detection of neural activity in functional mri using canonical correlation analysis. Magnetic Resonance in Medicine, 45(2):323–330, 2001. ISSN 1522-2594. doi: https://doi.org/10.1002/1522-2594(200102)45:2(323::AID-MRM1041)3.0.CO;2-#.

S. Gadgil, Q. Zhao, A. Pfefferbaum, E. V. Sullivan, E. Adeli, and K. M. Pohl. Spatiotemporal graph convolution for functional mri analysis, 2020.

Hélène Gervais, Pascal Belin, Nathalie Boddaert, Marion Leboyer, Arnaud Coez, Ignacio Sfaello, Catherine Barthélémy, Francis Brunelle, Yves Samson, and Mônica Zilbovicius. Abnormal cortical voice processing in autism. Nature Neuroscience, 7(88):801–802, 2004. ISSN 1546-1726. doi: 10.1038/nn1291.

Jack Goffinet, Richard Mooney, and John Pearson. Inferring low-dimensional latent descriptions of animal vocalizations. bioRxiv, page 811661, 2019. doi: 10.1101/811661.

K. A. N. N. P. Gunawardena, R. N. Rajapakse, and N. D. Kodikara. Applying convolutional neural networks for pre-detection of alzheimer’s disease from structural mri data. In 2017 24th International Conference on Mechatronics and Machine Vision in Practice (M2VIP), page 1–7, 2017. doi: 10.1109/M2VIP.2017.8211486.

A. N. Hampton and J. P. O’Doherty. Decoding the neural substrates of reward-related decision making with functional mri. Proceedings of the National Academy of Sciences, 104:1377–1382, 2007.

Xiaobing Han, Yanfei Zhong, Lifang He, Philip S Yu, and Liangpei Zhang. The unsupervised hierarchical convolutional sparse auto-encoder for neuroimaging data classification. page 12, 2015.

David R. Hardoon, Janaina Mourão-Miranda, Michael Brammer, and John Shawe-Taylor. Unsupervised analysis of fmri data using kernel canonical correlation. NeuroImage, 37 (4):1250–1259, 2007. ISSN 1053-8119. doi: 10.1016/j.neuroimage.2007.06.017.

Ryu-ichiro Hashimoto, KangUk Lee, Alexander Preus, Robert W. McCarley, and Cynthia G. Wible. An fmri study of functional abnormalities in the verbal working memory system and the relationship to clinical symptoms in chronic schizophrenia. Cerebral Cortex, 20(1):46–60, 04 2009.

Trevor J Hastie and Robert J Tibshirani. Generalized additive models, volume 43. CRC press, 1990.

L. Henschel, S. Conjeti, S. Estrada, K. Diers, B. Fischl, and M. Reuter. Fastsurfer - a fast and accurate deep learning based neuroimaging pipeline. NeuroImage, 219:117012, 2020.

James Hensman, Alexander Matthews, and Zoubin Ghahramani. Scalable variational gaussian process classification. In Artificial Intelligence and Statistics, pages 351–360. PMLR, 2015.

Irina Higgins, Loic Matthey, Arka Pal, Christopher Burgess, Xavier Glorot, Matthew Botvinick, Shakir Mohamed, and Alexander Lerchner. beta-vae: Learning basic visual concepts with a constrained variational framework. International Conference on Learning Representations, 2017.

Matthew D Hoffman and Matthew J Johnson. Elbo surgery: yet another way to carve up the variational evidence lower bound. In Workshop in Advances in Approximate Bayesian Inference, NIPS, volume 1, page 2, 2016.

H. Huang, X. Hu, J. Han, J. Lv, N. Liu, L. Guo, and T. Liu. Latent source mining in fmri data via deep neural network. In 2016 IEEE 13th International Symposium on Biomedical Imaging (ISBI), page 638–641, Apr 2016. doi: 10.1109/ISBI.2016.7493348.

Heng Huang, Xintao Hu, Yu Zhao, Milad Makkie, Qinglin Dong, Shijie Zhao, Lei Guo, and Tianming Liu. Modeling task fmri data via deep convolutional autoencoder. IEEE transactions on medical imaging, 37(7):1551–1561, 2017.

Hojin Jang, Sergey M. Plis, Vince D. Calhoun, and Jong-Hwan Lee. Task-specific feature extraction and classification of fmri volumes using a deep neural network initialized with a deep belief network: Evaluation using sensorimotor tasks. NeuroImage, 145:314–328, 2017. ISSN 1053-8119. doi: 10.1016/j.neuroimage.2016.04.003.

J.M Jansma, N.F Ramsey, N.J.A van der Wee, and R.S Kahn. Working memory capacity in schizophrenia: a parametric fmri study. Schizophrenia Research, 68(2):159 – 171, 2004.

Mark Jenkinson, Christian F. Beckmann, Timothy E. J. Behrens, Mark W. Woolrich, and Stephen M. Smith. Fsl. NeuroImage, 62(2):782–790, 2012. ISSN 1053-8119. doi: 10.1016/j.neuroimage.2011.09.015.

Huiting Jiang, Na Lu, Kewei Chen, Li Yao, Ke Li, Jiacai Zhang, and Xiaojuan Guo. Predicting brain age of healthy adults based on structural mri parcellation using convolutional neural networks. Frontiers in Neurology, 10, 2020. ISSN 1664-2295. doi: 10.3389/fneur.2019.01346. URL https://www.frontiersin.org/articles/10.3389/fneur.2019.01346/full.

B. A. Jonsson, G. Bjornsdottir, T. E. Thorgeirsson, L. M. Ellingsen, G. Bragi Walters, D. F. Gudbjartsson, H. Stefansson, K. Stefansson, and M. O. Ulfarsson. Brain age prediction using deep learning uncovers associated sequence variants. Nature Communications, 10 (11):5409, 2019. ISSN 2041-1723. doi: 10.1038/s41467-019-13163-9.

R. Ju, C. Hu, p. zhou, and Q. Li. Early diagnosis of alzheimer’s disease based on restingstate brain networks and deep learning. IEEE/ACM Transactions on Computational Biology and Bioinformatics, 16:244–257, 2019.

Ronghui Ju, Chenhui Hu, Pan Zhou, and Quanzheng Li. Early diagnosis of alzheimer’s disease based on resting-state brain networks and deep learning. IEEE/ACM Transactions on Computational Biology and Bioinformatics, 16(1):244–257, Jan 2019. ISSN 1557-9964. doi: 10.1109/TCBB.2017.2776910.

Meenakshi Khosla, Keith Jamison, Amy Kuceyeski, and Mert R. Sabuncu. Ensemble learning with 3d convolutional neural networks for functional connectome-based prediction. NeuroImage, 199:651–662, 2019. ISSN 1053-8119. doi: 10.1016/j.neuroimage.2019.06.012.

Diederik P Kingma and Max Welling. Auto-Encoding variational bayes. dec 2013.

Hideya Koshino, Patricia A. Carpenter, Nancy J. Minshew, Vladimir L. Cherkassky, Timothy A. Keller, and Marcel Adam Just. Functional connectivity in an fmri working memory task in high-functioning autism. NeuroImage, 24(3):810 – 821, 2005.

Tuan Anh Le, Maximilian Igl, Tom Rainforth, Tom Jin, and Frank Wood. Auto-encoding sequential monte carlo. arXiv preprint arXiv:1705.10306, 2017.

K. Lee, S. Tak, and J. C. Ye. A data-driven sparse glm for fmri analysis using sparse dictionary learning with mdl criterion. IEEE Transactions on Medical Imaging, 30(5): 1076–1089, 2011. ISSN 1558-254X. doi: 10.1109/TMI.2010.2097275.

X. Li, N. C. Dvornek, X. Papademetris, J. Zhuang, L. H. Staib, P. Ventola, and J. S. Duncan. 2-channel convolutional 3d deep neural network (2cc3d) for fmri analysis: Asd classification and feature learning. In 2018 IEEE 15th International Symposium on Biomedical Imaging (ISBI 2018), page 1252–1255, 2018. doi: 10.1109/ISBI.2018.8363798.

Dongdong Lin, Vince D. Calhoun, and Yu-Ping Wang. Correspondence between fmri and snp data by group sparse canonical correlation analysis. Medical Image Analysis, 18(6): 891–902, 2014. ISSN 1361-8415. doi: 10.1016/j.media.2013.10.010.

Jinglei Lv, Xi Jiang, Xiang Li, Dajiang Zhu, Hanbo Chen, Tuo Zhang, Shu Zhang, Xintao Hu, Junwei Han, Heng Huang, and et al. Sparse representation of whole-brain fmri signals for identification of functional networks. Medical Image Analysis, 20(1):112–134, 2015. ISSN 1361-8415. doi: 10.1016/j.media.2014.10.011.

Chris J Maddison, John Lawson, George Tucker, Nicolas Heess, Mohammad Norouzi, Andriy Mnih, Arnaud Doucet, and Yee Teh. Filtering variational objectives. In Advances in Neural Information Processing Systems, pages 6573–6583, 2017.

Abdelhak Mahmoudi, Sylvain Takerkart, Fakhita Regragui, Driss Boussaoud, and Andrea Brovelli. Multivoxel pattern analysis for fmri data: a review. Computational and mathematical methods in medicine, 2012, 2012.

Zhenyu Mao, Yi Su, Guangquan Xu, Xueping Wang, Yu Huang, Weihua Yue, Li Sun, and Naixue Xiong. Spatio-temporal deep learning method for adhd fmri classification. Information Sciences, 499:1–11, 2019. ISSN 0020-0255. doi: 10.1016/j.ins.2019.05.043.

T. Matsubara, K. Kusano, T. Tashiro, K. Ukai, and K. Uehara. Deep generative model of individual variability in fmri images of psychiatric patients. IEEE Transactions on Biomedical Engineering, 68(2):592–605, 2021. ISSN 1558-2531. doi: 10.1109/TBME.2020.3008707.

Takashi Matsubara, Tetsuo Tashiro, and Kuniaki Uehara. Structured deep generative model of fmri signals for mental disorder diagnosis. In Alejandro F. Frangi, Julia A. Schnabel, Christos Davatzikos, Carlos Alberola-López, and Gabor Fichtinger, editors, Medical Image Computing and Computer Assisted Intervention – MICCAI 2018, Lecture Notes in Computer Science, page 258–266. Springer International Publishing, 2018. ISBN 978-3-030-00931-1. doi: 10.1007/978-3-030-00931-1_30.

Amalia R. McDonald, Jordan Muraskin, Nicholas T. Van Dam, Caroline Froehlich, Benjamin Puccio, John Pellman, Clemens C.C. Bauer, Alexis Akeyson, Melissa M. Breland, Vince D. Calhoun, Steven Carter, Tiffany P. Chang, Chelsea Gessner, Alyssa Gianonne, Steven Giavasis, Jamie Glass, Steven Homann, Margaret King, Melissa Kramer, Drew Landis, Alexis Lieval, Jonathan Lisinski, Anna Mackay-Brandt, Brittny Miller, Laura Panek, Hayley Reed, Christine Santiago, Eszter Schoell, Richard Sinnig, Melissa Sital, Elise Taverna, Russell Tobe, Kristin Trautman, Betty Varghese, Lauren Walden, Runtang Wang, Abigail B. Waters, Dylan C. Wood, F.Xavier Castellanos, Bennett Leventhal, Stanley J. Colcombe, Stephen LaConte, Michael P. Milham, and R. Cameron Craddock. The real-time fmri neurofeedback based stratification of default network regulation neuroimaging data repository. NeuroImage, 146:157 – 170, 2017.

Christian Naesseth, Scott Linderman, Rajesh Ranganath, and David Blei. Variational sequential monte carlo. In International Conference on Artificial Intelligence and Statistics, pages 968–977. PMLR, 2018.

K.K. Nicodemus, J.H. Callicott, R.G. Higier, A. Luna, D.C. Nixon, B.K. Lipska, R. Vakkalanka, I. Giegling, D. Rujescu, D. St. Clair, P. Muglia, Y. Shugart, and D. Weinberger. Evidence of statistical epistasis between disc1, cit and ndel1 impacting risk for schizophrenia: biological validation with functional neuroimaging. Human Genetics, 127: 441–452, 2010.

Kate Nooner, Stanley Colcombe, Russell Tobe, Maarten Mennes, Melissa Benedict, Alexis Moreno, Laura Panek, Shaquanna Brown, Stephen Zavitz, Qingyang Li, Sharad Sikka, David Gutman, Saroja Bangaru, Rochelle Tziona Schlachter, Stephanie Kamiel, Ayesha Anwar, Caitlin Hinz, Michelle Kaplan, Anna Rachlin, Samantha Adelsberg, Brian Cheung, Ranjit Khanuja, Chaogan Yan, Cameron Craddock, Vincent Calhoun, William Courtney, Margaret King, Dylan Wood, Christine Cox, Clare Kelly, Adriana DiMartino, Eva Petkova, Philip Reiss, Nancy Duan, Dawn Thompsen, Bharat Biswal, Barbara Coffey, Matthew Hoptman, Daniel Javitt, Nunzio Pomara, John Sidtis, Harold Koplewicz, Francisco Castellanos, Bennett Leventhal, and Michael Milham. The nki-rockland sample: A model for accelerating the pace of discovery science in psychiatry. Frontiers in Neuroscience, 6:152, 2012.

Kenneth A Norman, Sean M Polyn, Greg J Detre, and James V Haxby. Beyond mindreading: multi-voxel pattern analysis of fmri data. Trends in cognitive sciences, 10(9): 424–430, 2006.

S. Ogawa, D. W. Tank, R. Menon, J. M. Ellermann, S. G. Kim, H. Merkle, and K. Ugurbil. Intrinsic signal changes accompanying sensory stimulation: Functional brain mapping with magnetic resonance imaging. Proceedings of the National Academy of Sciences, 89: 5951–5955, 1992.

William Penny, Karl Friston, John Ashburner, Stefan Kiebel, and Thomas Nichols. Statistical Parametric Mapping: The Analysis of Functional Brain Images. Elsevier Science & Technology, 2006.

Ruth C. M. Philip, Maria R. Dauvermann, Heather C. Whalley, Katie Baynham, Stephen M. Lawrie, and Andrew C. Stanfield. A systematic review and meta-analysis of the fmri investigation of autism spectrum disorders. Neuroscience & Biobehavioral Reviews, 36 (2):901–942, 2012. ISSN 0149-7634. doi: 10.1016/j.neubiorev.2011.10.008.

Ning Qiang, Qinglin Dong, Wei Zhang, Bao Ge, Fangfei Ge, Hongtao Liang, Yifei Sun, Jie Gao, and Tianming Liu. Modeling task-based fmri data via deep belief network with neural architecture search. Computerized Medical Imaging and Graphics, page 101747, 2020.

J.D. Ragland, B.I. Turetsky, R.C. Gur, F. Gunning-Dixon, T. Turner, L. Schroeder, R. Chan, and R.E. Gur. Working memory for complex figures: An fmri comparison of letter and fractal n-back tasks. Neuropsychology, 16(3):370–379, 2002.

Dehua Ren, Yu Zhao, Hanbo Chen, Qinglin Dong, Jinglei Lv, and Tianming Liu. 3-d functional brain network classification using convolutional neural networks. page 1217–1221, Apr 2017. doi: 10.1109/ISBI.2017.7950736.

Danilo Jimenez Rezende, Shakir Mohamed, and Daan Wierstra. Stochastic backpropagation and approximate inference in deep generative models. volume 32 of Proceedings of Machine Learning Research, pages 1278–1286, Bejing, China, 22–24 Jun 2014. PMLR. URL http://proceedings.mlr.press/v32/rezende14.html.

A. Riaz, M. Asad, S. M. M. R. A. Arif, E. Alonso, D. Dima, P. Corr, and G. Slabaugh. Deep fmri: An end-to-end deep network for classification of fmri data. In 2018 IEEE 15th International Symposium on Biomedical Imaging (ISBI 2018), page 1419–1422, 2018. doi: 10.1109/ISBI.2018.8363838.

Saman Sarraf and Ghassem Tofighi. Classification of alzheimer’s disease using fmri data and deep learning convolutional neural networks. arXiv:1603.08631 [cs], 2016. URL http://arxiv.org/abs/1603.08631.

Stephen M. Smith, Mark Jenkinson, Mark W. Woolrich, Christian F. Beckmann, Timothy E. J. Behrens, Heidi Johansen-Berg, Peter R. Bannister, Marilena De Luca, Ivana Drobnjak, David E. Flitney, and et al. Advances in functional and structural mr image analysis and implementation as fsl. NeuroImage, 23 Suppl 1:S208–219, 2004. ISSN 1053-8119. doi: 10.1016/j.neuroimage.2004.07.051.

H. Suk, C. Wee, S. Lee, and D. Shen. State-space model with deep learning for functional dynamics estimation in resting-state fmri. NeuroImage, 129:292–307, 2016.

Heung-Il Suk, Seong-Whan Lee, Dinggang Shen, and The Alzheimer’s Disease Neuroimaging Initiative. Latent feature representation with stacked auto-encoder for ad/mci diagnosis. Brain Structure and Function, 220(2):841–859, Mar 2015. ISSN 1863-2661. doi: 10.1007/s00429-013-0687-3.

Amirhessam Tahmassebi, Amir H. Gandomi, Ian McCann, Mieke H. J. Schulte, Anna E. Goudriaan, and Anke Meyer-Baese. Deep learning in medical imaging: fmri big data analysis via convolutional neural networks. In Proceedings of the Practice and Experience on Advanced Research Computing, PEARC ’18, page 1–4. Association for Computing Machinery, 2018. ISBN 978-1-4503-6446-1. doi: 10.1145/3219104.3229250. URL https://doi.org/10.1145/3219104.3229250.

Qiyuan Tian, Berkin Bilgic, Qiuyun Fan, Congyu Liao, Chanon Ngamsombat, Yuxin Hu, Thomas Witzel, Kawin Setsompop, Jonathan R. Polimeni, and Susie Y. Huang. Deepdti: High-fidelity six-direction diffusion tensor imaging using deep learning. NeuroImage, 219: 117017, 2020. ISSN 1053-8119. doi: 10.1016/j.neuroimage.2020.117017.

Christopher KI Williams and Carl Edward Rasmussen. Gaussian processes for machine learning, volume 2. MIT press Cambridge, MA, 2006.

B. C. Wittmann, B.H. Schott, S. Guderian, J. U. Frey, H. Heinze, and E. Düzel. Reward-related fmri activation of dopaminergic midbrain is associated with enhanced hippocampus-dependent long-term memory formation. Neuron, 45:459–467, 2005.

Mark W Woolrich, Timothy EJ Behrens, Christian F Beckmann, Mark Jenkinson, and Stephen M Smith. Multilevel linear modelling for fmri group analysis using bayesian inference. Neuroimage, 21(4):1732–1747, 2004.

Mark W. Woolrich, Saad Jbabdi, Brian Patenaude, Michael Chappell, Salima Makni, Timothy Behrens, Christian Beckmann, Mark Jenkinson, and Stephen M. Smith. Bayesian analysis of neuroimaging data in fsl. NeuroImage, 45(1 Suppl):S173–186, 2009. ISSN 1095-9572. doi: 10.1016/j.neuroimage.2008.10.055.

Zhengshi Yang, Xiaowei Zhuang, Karthik Sreenivasan, Virendra Mishra, Tim Curran, and Dietmar Cordes. A robust deep neural network for denoising task-based fmri data: An application to working memory and episodic memory. Medical Image Analysis, 60:101622, 2020. ISSN 1361-8415. doi: https://doi.org/10.1016/j.media.2019.101622. URL http://www.sciencedirect.com/science/article/pii/S1361841519301586.

Mariam Zabihi, Seyed Mostafa Kia, Thomas Wolfers, Richard Dinga, Alberto Llera, Danilo Bzdok, Christian F. Beckmann, and Andre Marquand. Non-linearity matters: a deep learning solution to generalization of hidden brain patterns across population cohorts. bioRxiv, page 2021.03.10.434856, 2021. doi: 10.1101/2021.03.10.434856.

Jun Zhang. Inverse-consistent deep networks for unsupervised deformable image registration. airXiv:1809.03443 [cs], 2018. URL http://arxiv.org/abs/1809.03443. arXiv: 1809.03443.

R. Zhang, X. Wei, and K. Kay. Understanding multivariate brain activity: Evaluating the effect of voxelwise noise correlations on population codes in functional magnetic resonance imaging. PLOS Computational Biology, 16:e1008153, 2020.

Chongyue Zhao, Hongming Li, Zhicheng Jiao, Tianming Du, and Yong Fan. A 3d convolutional encapsulated long short-term memory (3dconv-lstm) model for denoising fmri data. In Anne L. Martel, Purang Abolmaesumi, Danail Stoyanov, Diana Mateus, Maria A. Zuluaga, S. Kevin Zhou, Daniel Racoceanu, and Leo Joskowicz, editors, Medical Image Computing and Computer Assisted Intervention – MICCAI 2020, Lecture Notes in Computer Science, page 479–488. Springer International Publishing, 2020. ISBN 978-3-030-59728-3. doi: 10.1007/978-3-030-59728-3_47.

Qingyu Zhao, Ehsan Adeli, Nicolas Honnorat, Tuo Leng, and Kilian M Pohl. Variational autoencoder for regression: Application to brain aging analysis. In International Conference on Medical Image Computing and Computer-Assisted Intervention, pages 823–831. Springer, 2019.

Jacco A. de Zwart, Peter van Gelderen, Masaki Fukunaga, and Jeff H. Duyn. Reducing correlated noise in fmri data. Magnetic Resonance in Medicine, 59(4):939–945, 2008. ISSN 1522-2594. doi: https://doi.org/10.1002/mrm.21507.

