## Supplemental Materials for "Deep Generative Analysis for Task-Based Functional MRI Experiments"

### Appendix A. Network Architecture Details

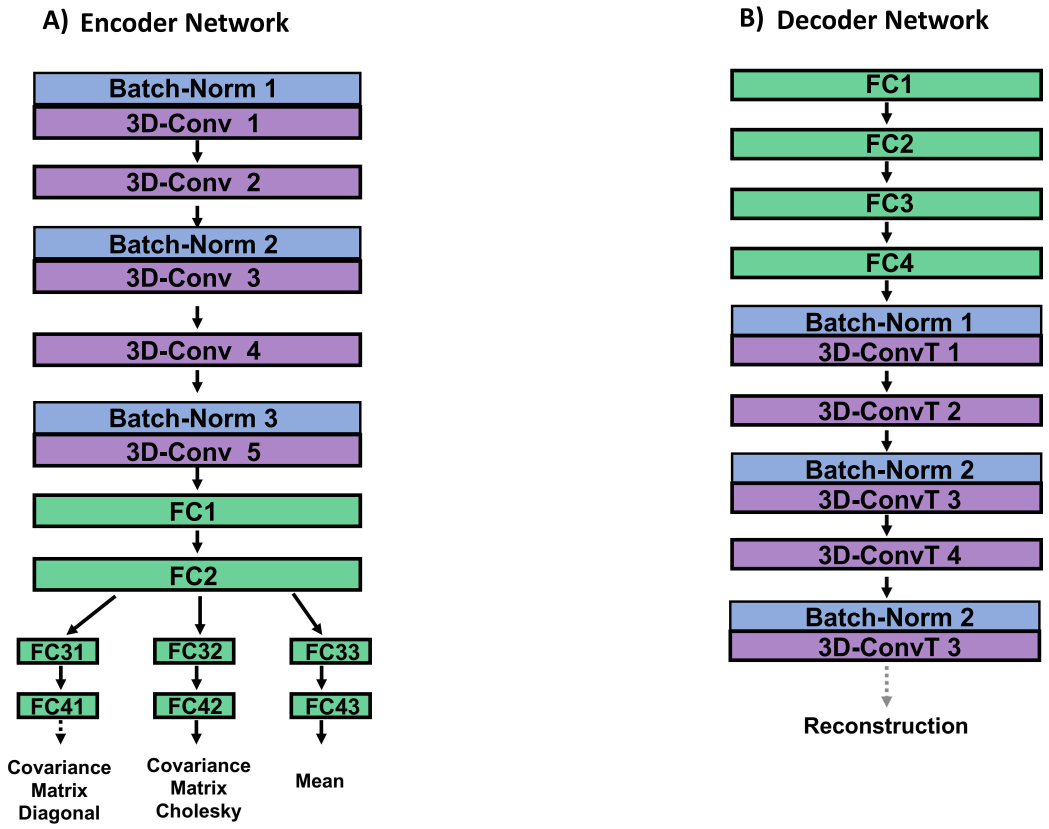


Figure 6: **Neural Network Architecture. Panel A:** Encoder network architecture. All 3D-convolutional layers have kernel size 3 and stride of either 1 (layers 1, 3 and 5) or 2 (layers 2 and 4). **Panel B:** Decoder network architecture. Transposed 3D-convolutional layers 1, 3 and 5 have kernel size 3 and stride 1. Transposed 3D-conv layer 2 has kernel size 3 and stride 2. Finally, transposed 3D-convolutional layer 4 has kernel size (5, 3, 3) and stride of 2. Solid black arrows indicate ReLU activation functions. The dashed black arrow indicates an exponential function (to enforce positivity of variances) and the dashed gray arrow a sigmoid function.

Table 1: Encoder Network Parameter Counts and Shapes

| Layer | Weights | Bias | Shape |
|---|---|---|---|
| Conv_1 | 216 | 8 | [8, 1, 3, 3, 3] |
| Batch_norm_1 | 1 | 1 | 1 |
| Conv_2 | 1728 | 8 | [8, 8, 3, 3, 3] |
| Conv_3 | 3456 | 16 | [16, 8, 3, 3, 3] |
| Batch_norm_3 | 8 | 8 | 8 |
| Conv_4 | 6912 | 16 | [16, 16, 3, 3, 3] |
| Conv_5 | 6912 | 16 | [16, 16, 3, 3, 3] |
| Batch_norm_5 | 16 | 16 | 16 |
| Fc_1 | 1614400 | 200 | [200, 3072] |
| Fc_2 | 20000 | 100 | [100, 200] |
| Fc_31, Fc_32, Fc_33 | 5000 | 50 | [50, 100] |
| Fc_41, Fc_42, Fc_43 | 1600 | 32 | [32, 50] |

Table 2: Decoder Network Parameter Counts and Shapes

| Layer | Weights | Bias | Shape |
|---|---|---|---|
| Fc_5 | 2000 | 850 | [50, 40] |
| Fc_6 | 5000 | 100 | [100, 50] |
| Fc_7 | 20000 | 200 | [200, 100] |
| Fc_8 | 768000 | 3840 | [3840, 200] |
| Convt_1 | 6912 | 16 | [16, 16, 3, 3, 3] |
| Batch_normt_1 | 16 | 16 | 16 |
| Convt_2 | 6912 | 16 | [16, 16, 3, 3, 3] |
| Convt_3 | 3456 | 8 | [16, 8, 3, 3, 3] |
| Batch_normt_3 | 16 | 16 | 16 |
| Convt_4 | 2880 | 8 | [8, 8, 5, 3, 3] |
| Convt_5 | 216 | 1 | [8, 1, 3, 3, 3] |
| Batch_normt_5 | 8 | 8 | 8 |

### Appendix B. Sparse Gaussian Process Details

Here, we clarify the derivation of our approximate GP objective (5) and the total objective (6). For the approximate posterior, we assume $q(\mathbf{z}, \boldsymbol{\beta}|\mathbf{x}) = q(\boldsymbol{\beta}|\mathbf{x}) \prod_t q(\mathbf{z}_t|\mathbf{x}_t)$ and write the full ELBO as

$$\begin{aligned}\log p(\mathbf{x}) &\geq \sum_t \mathbb{E}_q\left[\log p(\mathbf{x}_t|\mathbf{z}, \boldsymbol{\beta})\right] + \mathbb{E}_q[\log p(\mathbf{z}) + \log p(\boldsymbol{\beta})] + \mathcal{H}[q] \\ &= \sum_t \mathbb{E}_{q(\boldsymbol{\beta}|\mathbf{x}_t)} \mathbb{E}_{q(\mathbf{z}_t|\mathbf{x}_t)} \log p(\mathbf{x}_t|\mathbf{z}_t, \boldsymbol{\beta}) - D_{KL}(q(\mathbf{z}) \| p(\mathbf{z})) - D_{KL}(q(\boldsymbol{\beta}) \| p(\boldsymbol{\beta})) \\ &\approx \text{ELBO}(\bar{\boldsymbol{\beta}}) - D_{KL}(q(\boldsymbol{\beta}) \| p(\boldsymbol{\beta}))\end{aligned} \tag{7}$$

where ELBO denotes the evidence lower bound due to $\mathbf{z}$ alone, and we have approximated the expectation over the GPs $\boldsymbol{\beta}$ by using a plug-in estimator: their predicted means at the locations of the observed minibatch data, $\bar{\boldsymbol{\beta}}$. The remaining term in (7) can be recognized as a regularizer that encourages the approximate posterior $q(\boldsymbol{\beta})$ to remain close to its prior (Hoffman and Johnson, 2016), and it is well-established that there are benefits to both disentangling in the latent space and robustness in upweighting such terms (Higgins et al., 2017; Hoffman and Johnson, 2016). We thus weight this term by an additional hyperparameter $\gamma$ in (6).

Finally, for the form of the KL divergence between the GP prior and posterior, we follow (Hensman et al., 2015) in writing (for a single, univariate GP)

$$q(\mathbf{f}) = \int p(\mathbf{f}|\mathbf{u}) q(\mathbf{u}) d\mathbf{u}, \tag{8}$$

where $\mathbf{f} \equiv \delta\beta(\mathbf{c})$ is the vector of GP values evaluated at the data points and $q(\mathbf{u}) = \mathcal{N}(\mathbf{m}, \mathbf{S})$ is a variational Gaussian posterior for the values of the GP at a set of unknown inducing points $\mathbf{Z}$ (distinct from the latent variables $\mathbf{z}$ above). In the sparse variational GP approximation, the locations $\mathbf{Z}$ and GP values $\mathbf{u}$ of the inducing points are variational parameters to be optimized over, along with the parameters of the kernel $k(\cdot, \cdot)$. Moreover, $q(\mathbf{f})$ can be calculated in closed form (Hensman et al., 2015). For this work, we simplified this setup by choosing a fixed number of equally spaced inducing points over the observed range of each covariate. That is, we optimized over $\mathbf{u}$ but not $\mathbf{Z}$. In addition, since we only made use of the mean GP prediction $\overline{\delta\boldsymbol{\beta}} = \mathbf{K}_{nu}\mathbf{K}_u^{-1}\mathbf{m}$ in (7), this is equivalent to treating $\mathbf{f}$ as a deterministic function of $\mathbf{u}$. In this case, we have

$$\begin{aligned}D_{KL}(q(\boldsymbol{\beta}) \| p(\boldsymbol{\beta})) &\approx D_{KL}(q(\mathbf{u}) \| p(\mathbf{u})) \\ &= \frac{1}{2}\left(\mathbf{m}^\top \mathbf{K}_u^{-1}\mathbf{m} + \mathrm{tr}(\mathbf{K}_u^{-1}\mathbf{S}) + \log|\mathbf{K}_u| - \log|\mathbf{S}| - M\right),\end{aligned} \tag{9}$$

where $\mathbf{K}_u$ is the matrix formed by evaluating the kernel at the locations of the $M$ inducing points.

Alternatively, for our experiments, we make use of an maximum likelihood approach in which we do not perform inference on $\boldsymbol{\beta}$ using $q(\boldsymbol{\beta})$ but simply let $p(\mathbf{u}) = \mathcal{N}(0, \mathbf{K}_u)$ and learn both $\mathbf{u}$ and the parameters $\ell_\alpha$ and $\sigma_\alpha^2$ of the RBF kernels $k_\alpha(x, x') = \sigma_\alpha^2 \exp\left(-\frac{1}{2\ell_\alpha^2}(x - x')^2\right)$ that maximize

$$\log p(\mathbf{u}) = -\sum_\alpha \frac{1}{2}\left(\mathbf{u}_\alpha^\top \mathbf{K}_{u\alpha}^{-1}\mathbf{u}_\alpha + \log|\mathbf{K}_{u\alpha}| + M\log(2\pi)\right), \tag{10}$$

which is equivalent to (5).

### Appendix C. Simulations Performed with Different Hyper-Parameter Values

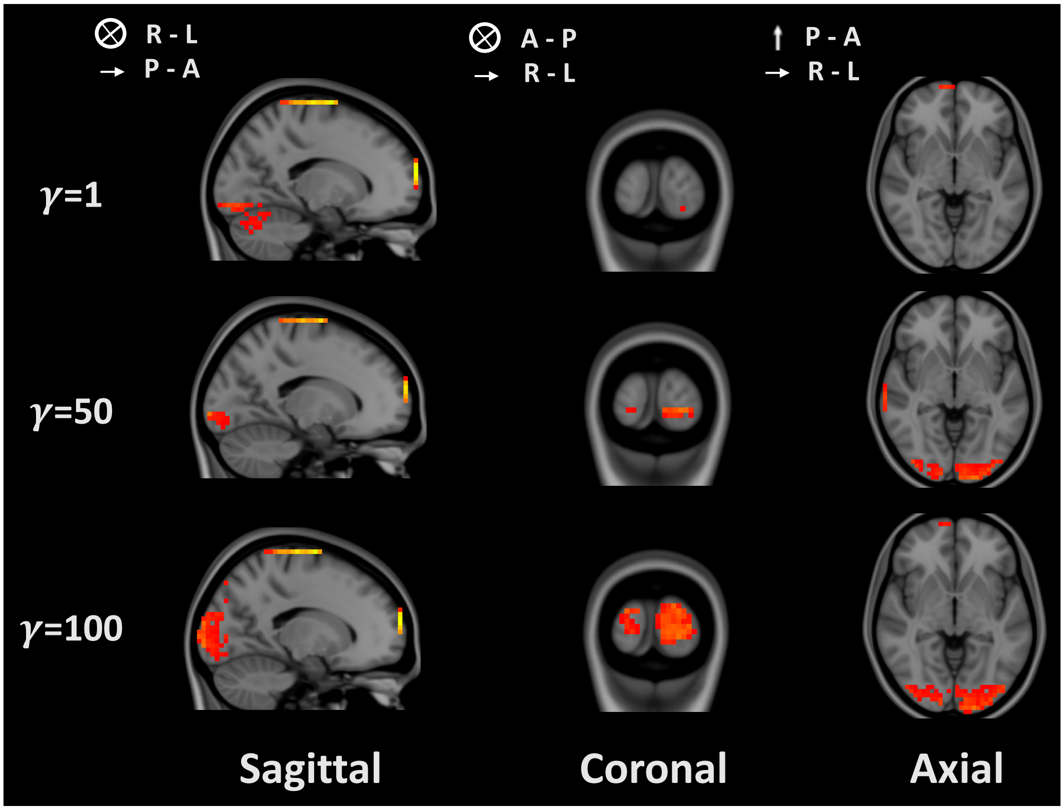


Figure 7: **Sensitivity to hyperparameter $\gamma$.** Inferred visual task effects for $\gamma \in \{1, 50, 100\}$ with $\delta = 0.005$. For lower values of $\gamma$, where the GP contribution to overall objective is small, the resulting maps do not properly capture either the spatial location or extent of the effect. At higher $\gamma$, the GP contribution to the objective increases, and model is better able to appropriately capture the location and size of the main effect cluster.

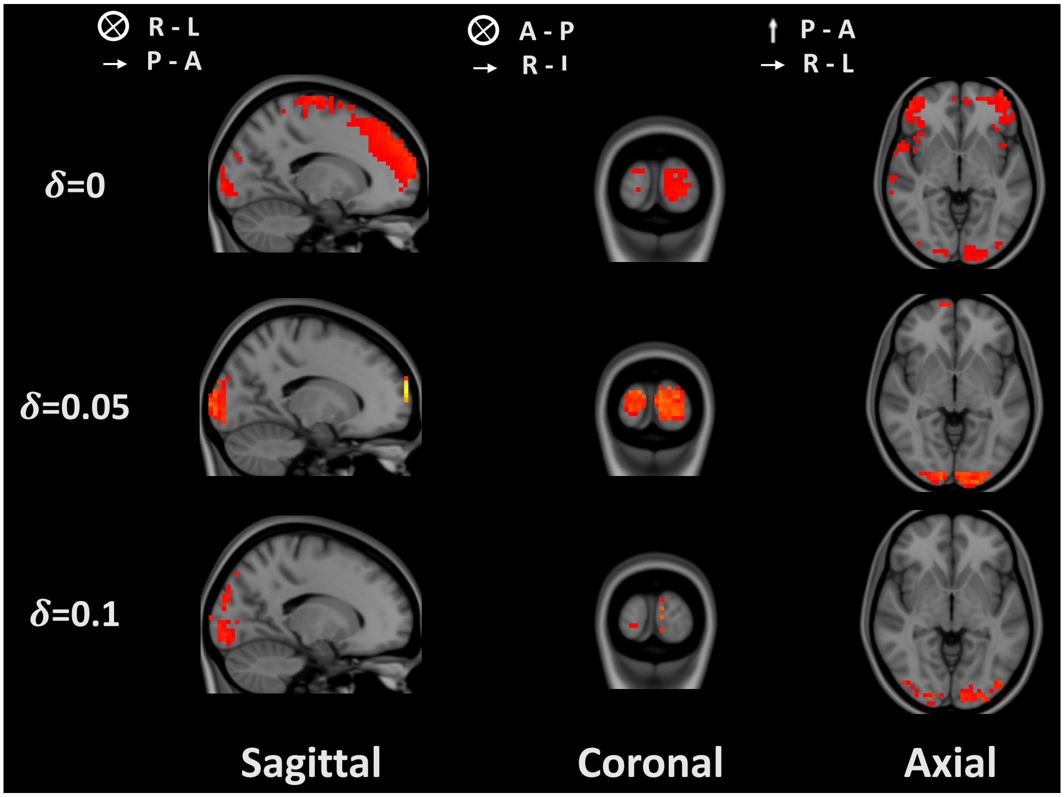


Figure 8: **Sensitivity to hyperparameter $\delta$.** Inferred visual task effects for $\delta \in \{0, 0.05, 0.1\}$ with $\gamma = 10$. As the experiment shows, levels of $L_1$ regularization of the spatial maps remove spurious activations in the frontal lobe, while higher levels of regularization reduce the inferred maps to extremely sparse microclusters of voxels. Nonetheless, these remaining voxels still lie within the appropriate regions.

### Appendix D. Additional Individual Subject Map Samples

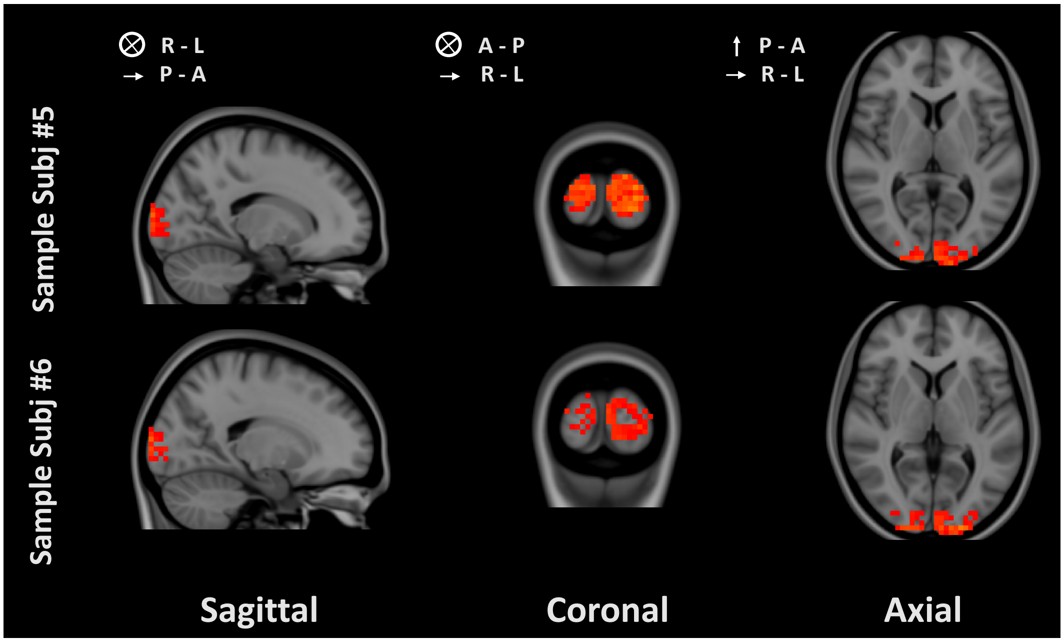


Figure 9: **Single subject average task maps for remaining 2 participants in our cohort.** Conventions are as in Figure 5. As before, note individual differences in the exact location and spatial extent of the inferred task effect.
